# Adaptations in the echolocation behavior of fruit-eating bats when orienting under challenging conditions

**DOI:** 10.1101/485490

**Authors:** M. Jerome Beetz, Manfred Kössl, Julio C. Hechavarría

## Abstract

**Summary statement:** When echolocating under demanding conditions e.g. noisy, narrow space, or cluttered environments, frugivorous bats adapt their call pattern by increasing the call rate within biosonar groups.

**Abstract:** For orientation, echolocating bats emit biosonar calls and use echoes arising from call reflections. They often pattern their calls into groups which increases the rate of sensory feedback over time. Insectivorous bats emit call groups at a higher rate when orienting in cluttered compared to uncluttered environments. Frugivorous bats increase the rate of call group emission when they echolocate in noisy environments. Here, calls emitted by conspecifics potentially interfere with the bat’s biosonar signals and complicate the echolocation behavior. To minimize the information loss followed by signal interference, bats may profit from a temporally increased sensory acquisition rate, as it is the case for the call groups. In frugivorous bats, it remains unclear if call group emission represents an exclusive adaptation to avoid interference by signals from other bats or if it represents an adaptation that allows to orient under demanding environmental conditions. Here, we compared the emission pattern of the frugivorous bat *Carollia perspicillata* when the bats were flying in noisy versus silent, narrow versus wide or cluttered versus non-cluttered corridors. According to our results, the bats emitted larger call groups and they increased the call rate within the call groups when navigating in narrow, cluttered, or noisy environments. Thus, call group emission represents an adaptive behavior when the bats orient in complex environments.

## Introduction

For collision-free locomotion, animals constantly update the location of surrounding obstacles. With increasing obstacle density, successful orientation becomes challenging and some animals increase their sensory acquisition rate (Geva-Sagiv et al., 2015). Electrolocating and echolocating animals utilize self-emitted signals and echoes for orientation (Geva-Sagiv et al., 2015; Hofmann et al., 2013; Kössl et al., 2014; Moss and Surlykke, 2010; Nelson and MacIver, 2006; Neuweiler, 1990). The amount of signals emitted within a certain period represent the sensory acquisition rate. This makes the acquisition rate highly accessible and allows us to answer questions on how bats adapt their sensory acquisition rate when orienting under different conditions. Many behavioral studies showed that bats often pattern their echolocation calls in form of groups (Amichai et al., 2015; Brinklov et al., 2011; Brinklov et al., 2009; Galambos and Griffin, 1942; Grinnell and Griffin, 1958; Kothari et al., 2018a; Luo et al., 2015; Roverud and Grinnell, 1985a; Roverud and Grinnell, 1985b; Wheeler et al., 2016; Wohlgemuth et al., 2016; Figure 1). Results from insectivorous bats led to the hypothesis that the emission of call groups may represent an adaptation to orient in complex environments (Falk et al., 2014; Fawcett et al., 2015; Kothari et al., 2014; Moss et al., 2006; Petrites et al., 2009; Sändig et al., 2014; Surlykke et al., 2009). Frugivorous bats emit more and larger call groups when orienting in the presence than in the absence of acoustic playbacks (Beetz et al., 2018; Luo et al., 2015). Acoustic playbacks potentially interfere with the bat’s echolocation system making echolocation highly demanding. Thus, for frugivorous bats, it has been proposed that the call groups may represent an adaptation to avoid signal interference (Beetz et al., 2018). However, it remains unknown if frugivorous bats show similar adaptations when orienting in narrow-spaced or cluttered environments as it has been shown for insectivorous bats. To clarify the role of call group emissions in frugivorous bats, the present study characterizes the call emission pattern of *Carollia perspicillata*, when the bats were flying in narrow versus wide or in cluttered versus non-cluttered corridors. We hypothesized that if *C. perspicillata* shows similar adaptations as it was previously demonstrated with the playback experiments (Beetz et al., 2018), then the adaptations are not exclusive to avoid acoustic interference, but they rather assist echolocation under highly demanding conditions.

**Figure 1:**
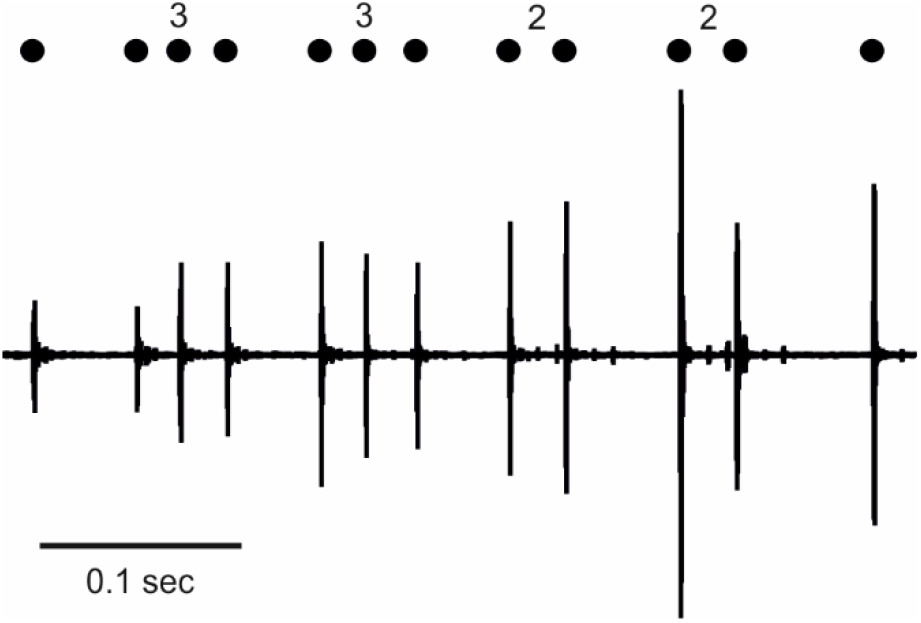
Echolocation sequence of *Carollia perspicillata* showing echolocation calls that are patterned into call groups. The oscillogram shows twelve echolocation calls of *C. perspicillata*.Black dots on top of the oscillogram signal the time points of call emission. The numbers on top of the dots represent the call group size (3 = triplet, 2 = doublet).

## Materials and methods

### Animals

Experiments were conducted in 45 bats of the species *Carollia perspicillata*. The bats were bred and kept in a colony at the Institute for Cell Biology and Neuroscience (Goethe-University Frankfurt). The experiments comply with all current German laws on animal experimentation and they are in accordance with the Declaration of Helsinki. All experimental protocols were approved by the Regierungspräsidium Darmstadt (experimental permit # FU-1126).

### Flight room

The experiments were performed in a flight room (length: 4 m; width: 1.4 m; height: 2 m). A wall, made out of foam, separated the room into two corridors. At the end of each corridor, a landing platform (20 x 20 cm), made out of metal mesh, was positioned. Behind each metal mesh, one speaker (Neo CD 1.0 Ribbon Tweeter; Fountek Electronics, China) and one ultrasound sensitive microphone (Avisoft Bioacoustics, Germany) were installed. The emission of the echolocation calls was monitored by the microphones which had a sensitivity of 50 mV/Pa and an input-referred self-noise level of 18 dB SPL. Each microphone was connected to a sound acquisition system (one microphone to an UltraSoundGate 116 Hm mobile recording interface and the second microphone to an UltraSoundGate 116 Hb mobile recording interface, + Recorder Software, Avisoft Bioacoustics, Germany) for sound digitalization at 333 kHz (16-bit precision). Bats were hand-released at one side of the flight room (starting position in figure 2) and they could freely fly in the flight room. The flight behavior was monitored with a webcam (500 SX, Manhattan, USA) placed above the starting point (frame rate = 30 Hz). The trial ended when the bat land on one of the two platforms. Since the ultrasound sensitive microphones were directly behind the landing platforms, the bats directly approached one of the microphones before ending a trial. This allows to record the patterns of the emitted echolocation calls while the bat was approaching the platform.

**Figure 2:**
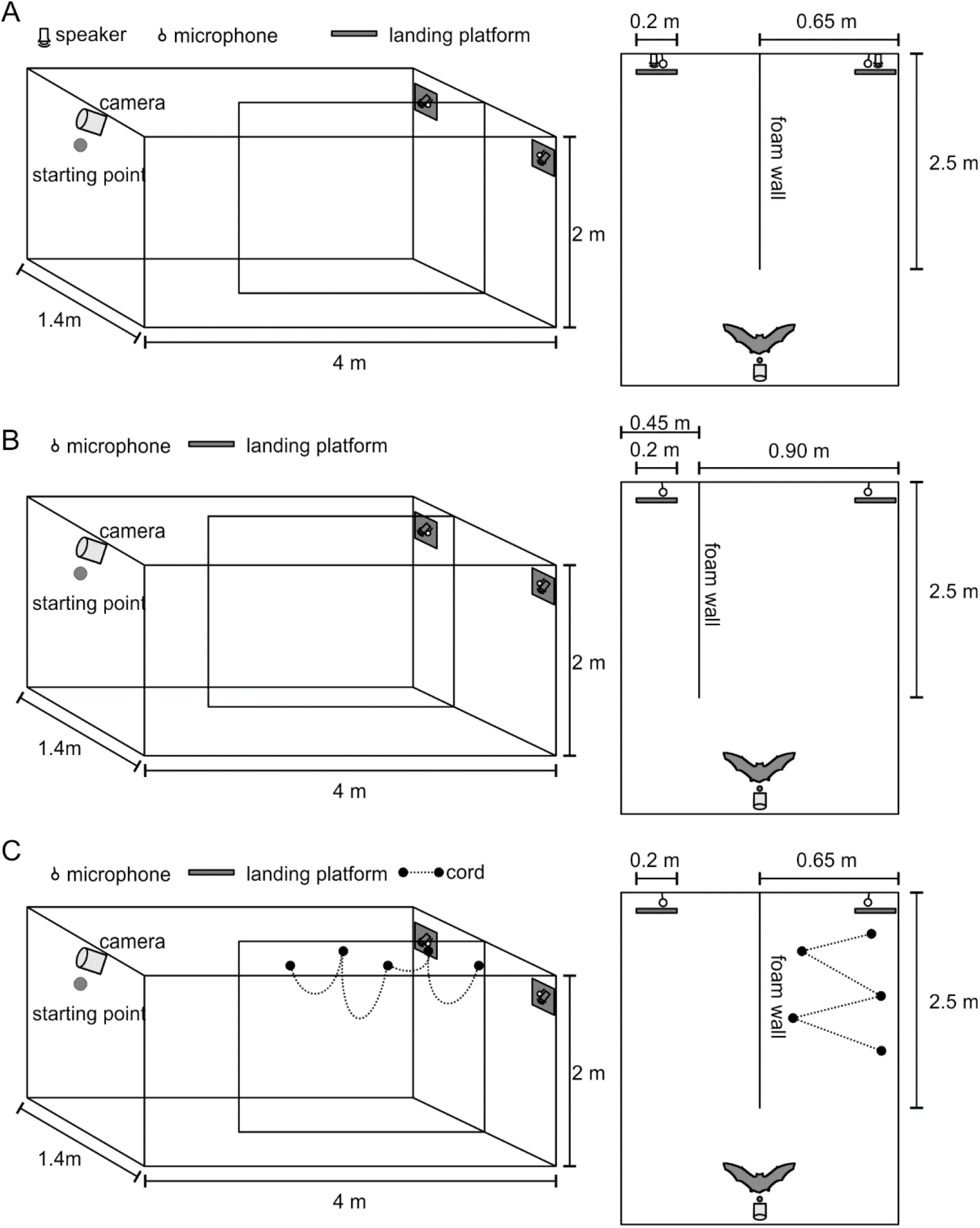
Schematic lateral and top views of the experimental designs for the three experiments. The experiments were conducted in a 1.4 m x 4 m x 2 m flight room. A moveable wall made of foam separated about two third of the room into two corridors. At the end of each corridor, the bats could land on one landing platform made of metal mesh. The bats were hand released at the starting point and they could freely fly in the flight room. A camera on top of the starting point and two ultrasound sensitive microphones, one positioned behind each landing platform could record the echolocation behavior of the animal. (**A**) In the first experiment, the speaker behind one of the landing platforms produced an acoustic playback making one corridor noisy (challenging corridor). The second corridor was silent (non-challenging corridor). (**B**) In the second experiment, the bats could either fly in a narrow (challenging) or in a wide (non-challenging) corridor. (**C**) In the third experiment, the bats could either fly in a cluttered (challenging) or in a non-cluttered (non-challenging) corridor. Clutter was represented, as loops of cord hanging from the ceiling of the flight room.

### Experiment 1: Influence of high frequency playback on call emission pattern

For comparative reasons, in the present report, data from a previously published manuscript (Beetz et al., 2018) are used. The echolocation behavior from eight bats was tested in experiment 1. We investigated the influence of acoustic playbacks containing high frequency echolocation calls on the echolocation behavior by presenting the bats one noisy and one silent corridor (Figure 2A). The call emission pattern emitted by the bat while flying in the noisy corridor (test trial) was compared with the emission pattern as the bat flew under entirely silent conditions (training trial). The playback stimuli represented repetitions of representative echolocation calls emitted by the tested bat during the training trials. The played back echolocation calls were repeated in groups of five, ten, or twenty calls. The call rate within the call groups was 66 Hz and the groups were repeated every 35 ms. Acoustic stimuli were generated with a sampling rate of 384 kHz with an Exasound E18 sound card (ExaSound Audio Design, Canada), and sent to an audio amplifier (Rotel power amplifier, RB-850, USA). The stimuli were played with a sound pressure level of 80-90 dB re 20 μPascal (dB SPL). For analysis, a sequence of two seconds from each trial was selected. Two seconds was usually the time window that the bats needed to approach and land on the platform for each corridor. During the approach flight the echolocation calls were intense enough to be easily detected by the microphone behind the platform. This allows use to ensure that we did not miss any echolocation call emitted during this sequence and that the recorded echolocation pattern represents the most “natural” one we could observe under these paradigm settings. In total, 48 sequences, six (three training and three test trials) from each animal were analyzed. Call emission pattern emitted during test and training trials were compared pairwise. Thus, three pairs of “test” and “training” trials were compared for each animal.

### Experiment 2: Influence of corridor width on call emission pattern

The influence of the corridor width on the echolocation behavior was tested in 21 bats. We modified the flight room so that the bats could choose flying in a narrow (0.45 m) or in a wide (0.9 m) corridor (Figure 2B). For each bat, we compared the echolocation behavior when flying in the narrow corridor with the behavior elicited when flying in the wide corridor. Thus, two sequences of the recording, each lasting 2 seconds, was selected for data analysis.

### Experiment 3: Influence of clutter on call emission pattern

The influence of clutter on the echolocation behavior was tested in 16 bats. The “cluttered” corridor was equipped with a cord that formed four diagonal loops hanging from the corridor’s ceiling (Figure 2C). The uncluttered corridor was free of cord. As in experiment 2, for each bat, a recording sequence was compared when flying in the cluttered corridor with a sequence recorded while the animal was flying in the uncluttered corridor. Each sequence lasted for 2 seconds.

### Analysis

For data analysis, the call emission time points were manually tagged in the software Avisoft SAS Lab Pro (Avisoft Bioacoustics, Germany). The rest of the analysis, except the statistical analysis, was done in a custom written script in Matlab 2014 (MathWorks, USA). Call groups were defined according to the criterions of (Kothari et al., 2014). A call group needs to be temporally isolated (“island criterion”). A temporal isolation is fulfilled, when the preceding and following call interval of a call group are 20% longer than the call intervals within a call group. The size of a call group, indicated by the number of calls of the call group, is defined by the “stability criterion”. For the fulfillment of the “stability criterion”, the call intervals within the call groups need to be invariant with 5% tolerance. Note that doublets, i.e. call groups containing two calls, can only fulfill the “island criterion”. For defining triplets, quartets, quintets, or sextets, both criteria need to be fulfilled.

For statistical analysis, we used the software GraphPad Prism 7 (GraphPad Software, USA; * p < 0.05; ** p < 0.01; *** p < 0.0001). Since the echolocation behavior in two conditions (control versus test trials) were compared to each other, statistical tests were either based on non-parametric Wilcoxon signed-rank test (W; in case of non-Gaussian distribution) or on parametric paired t-Tests (in case of Gaussian distribution).

## Results

We simulated three different scenarios, where the bats had to orient under highly demanding conditions (Figure 2). For the first experiment, we challenged the bats by presenting playbacks consisting of echolocation calls while the bats had to fly and echolocate in a flight room (Figure 2A). The playback stimulus represented a sequence of echolocation calls that was recorded initially from the tested animal. Since the playback stimuli were presented only in one of the two corridors, the bats could choose between a noisy or silent corridor (Figure 2A). The influence of acoustic playback on the echolocation behavior was tested in eight bats. Note that the behavioral results from the playback experiment have recently been published elsewhere (Beetz et al., 2018) and the results are described here only to compare the echolocation behavior across different scenarios. Not only acoustic signals which may interfere with the echolocation system make collision-free echolocation challenging but also the corridor width may affect the echolocation pattern. Thus, in the second experiment, we challenged the bats by narrowing (0.45 m) one and widening (0.9 m) the other corridor (Figure 2B). Under these conditions, 21 one bats were tested. For the third experiment, 16 bats oriented in a flight room that had a cluttered and a non-cluttered corridor (Figure 2C). Here, both corridors were equal in size but both differed by the presence or absence of clutter, represented by cord hanging as loops from the corridor’s ceiling. For all experiments, the bats had only two landing positions, one platform at the end of each corridor. Behind the platforms, ultrasound sensitive microphones recorded the bats’ call emission patterns. Representative echolocation sequences for each paradigm are presented in figure 3. As it can be noted in figure 3, the bats grouped their echolocation calls while flying in the flight room. This can be seen by looking at the time points of call emission, indicated as black dots on top of each sequence. The call group size and the call rate within the call groups, indicated by a reduced inter-call time interval, increased when the bats oriented in the more challenging corridor. This adaptation occurred irrespective of the nature of the challenge, i.e. during presenting acoustic playbacks (Figure 3A), narrowing the corridor (Figure 3B), or enriching the corridor with clutter (Figure 3C). The tendency of grouping the calls was always higher in the challenging (noisy, Figure 3A; narrow, Figure 3B; cluttered, Figure 3C) than in the non-challenging corridor (silent, Figure 3A; wide, Figure 3B; non-cluttered corridor, Figure 3C).

**Figure 3:**
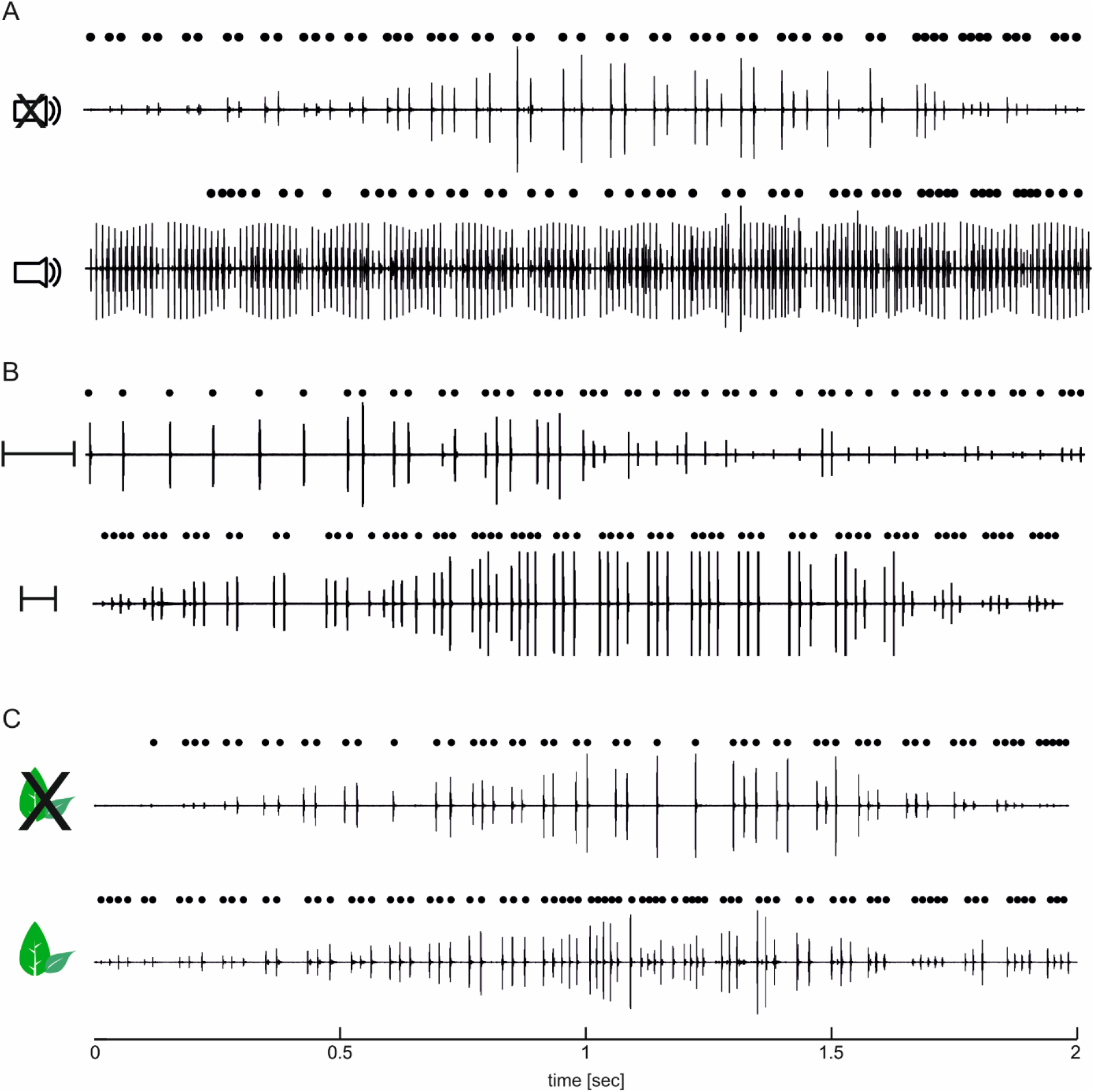
Representative echolocation sequences emitted in each corridor for each of the three experiments. Representative oscillograms visualizing the call emission pattern in a silent (upper graph in **A**), noisy (lower graph **A**), wide (upper graph **B**), narrow (lower graph **B**), uncluttered (upper graph **C**), and cluttered corridor (lower graph **C**). The time points of call emission are indicated as dots above each oscillogram. Note that the acoustic playback can also be seen as deflections in the oscillogram in (**A**). Under challenging conditions (lower graphs), the bats emitted more, larger, and more tightly packed call groups than under non-challenging conditions (upper graphs).

In the challenging corridor, the bats significantly reduced the minimum call-interval (Figure 4A). Note that the amount of reduction in minimum call-interval was comparable for each of the three experiments. This suggests that the reduction in minimum call-interval represents an adaptation to echolocate under demanding conditions rather than representing an exclusive adaptation to avoid only signal interference. The bats reduced the median call-interval only when orienting in the cluttered and narrowed corridor (Figure 4B). Acoustic playbacks had no significant effect on the median call-interval.

**Figure 4:**
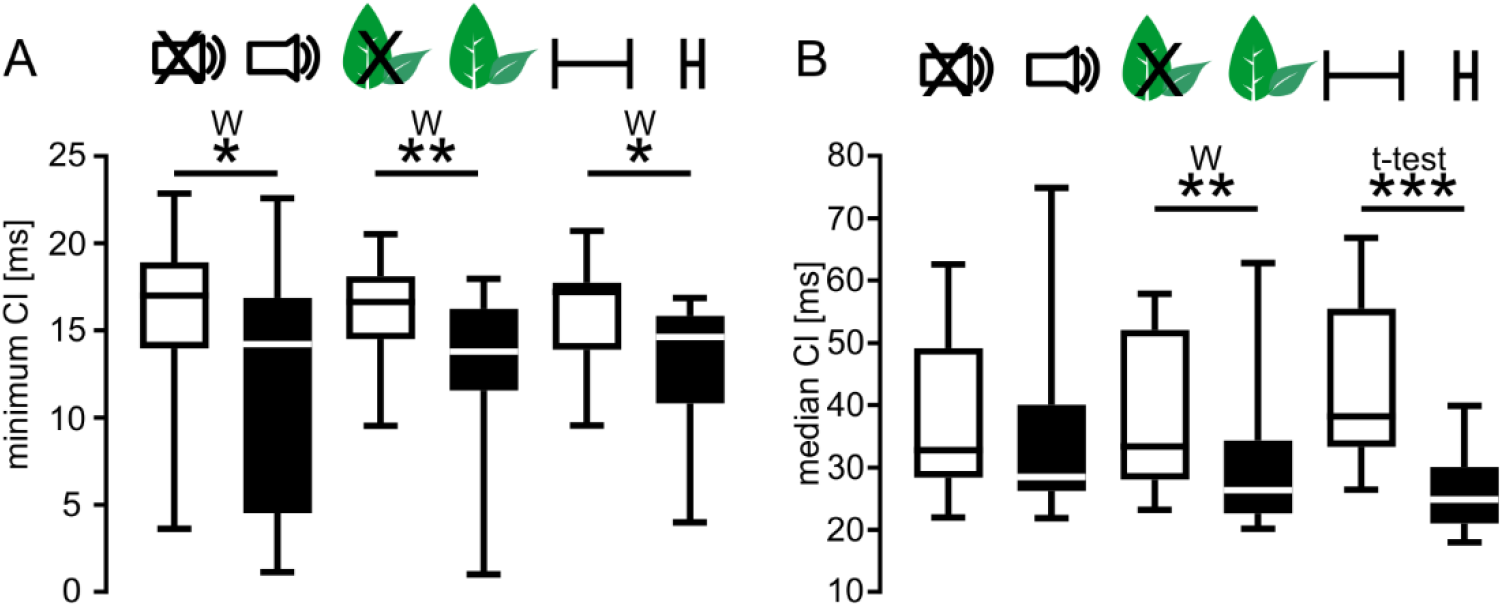
Boxplots showing the minimum and median call interval (CI) for each experimental condition. The bats decreased the minimum CI, under challenging conditions (black boxplots in A). They decreased the median CI when navigating in cluttered and narrow corridors as (black boxplots in B) as compared to non-cluttered and wide corridors (white boxplots in B). The presence of acoustic playback does not result into a significant decrease of the median CI as compared to the silent conditions. W = Wilcoxon signed rank test; t-Test = paired t-Test. * p < 0.05; ** p < 0.01; *** p < 0.0001

The relative amount of calls emitted as groups did not vary between the three experiments or between the challenging and non-challenging condition within each experiment (Figure 5A). About two thirds of the calls were emitted in form of call groups irrespective of the task or its complexity. In the challenging corridor, the bats reduced the call-intervals within the call groups resulting into a higher call rate within the call groups (Figure 5B). The extent of call rate increase within the call groups was comparable for each paradigm. This shows again that the call rate increase is not exclusive to avoid jamming. By taking a closer look into the call group size, indicated by the amount of calls per call group (two for doublet, three for triplet, four for quartet, five for quintet, and six for sextet), it becomes clear that the bats emitted significantly more triplets in the noisy than in the silent corridor (Figure 5C). In the cluttered and narrow corridors, the bats emitted significantly more quartets than in the non-cluttered and wide corridors (Figure 5D and 5E). Despite the difference in call group size across the three experiments, it is noteworthy, that the tendencies of emitting larger call groups in challenging than in non-challenging corridors was present in each of the three paradigms.

**Figure 5:**
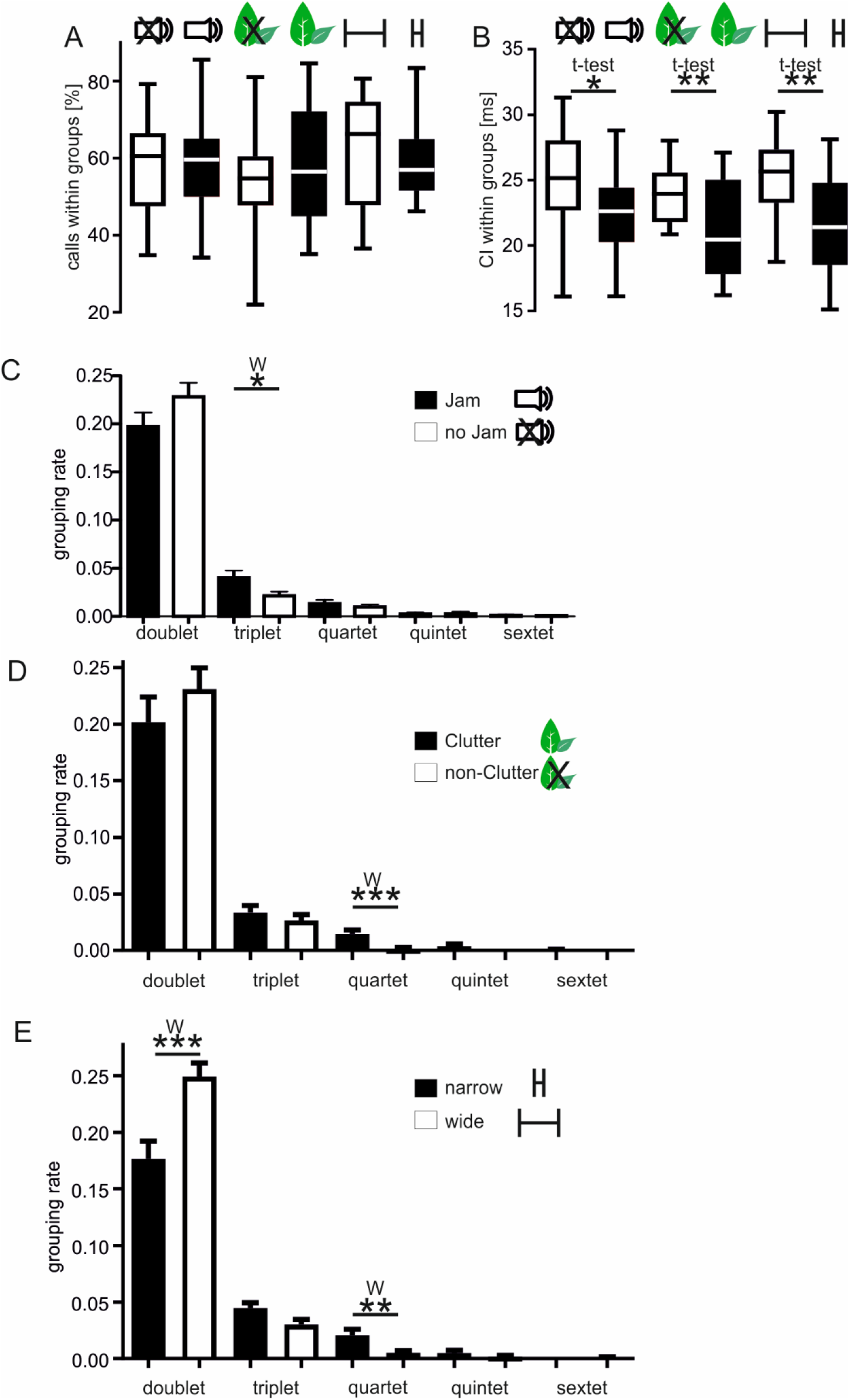
Parameters of the call groups. (**A**) When comparing the different experimental conditions, the bats did not change the relative amount of calls that were emitted as call groups. (**B**) When navigating under challenging conditions (noisy, narrow, or cluttered), the bats decrease the call intervals within the call groups as compared to the call intervals shown in less demanding conditions (silent, wide, or non-cluttered). (**C-E**) Histograms showing the relative amount of doublets, triplets, quartets, quintets, and sextets. (**C**) The bats emitted relatively more triplets when navigating in the noisy than in the silent corridor. (**D**) The bats emitted relatively more quartets when navigating in the cluttered than in the uncluttered corridor. (**E**) The bats emitted relatively more quartets and less doublets when navigating in the narrow than in the wide corridor. W = Wilcoxon signed rank test; t-Test = paired t-Test. * p < 0.05; ** p < 0.01; *** p < 0.0001

## Discussion

Animals often orient in habitats that are enriched with many obstacles. Under such conditions, rapid and collision-free movements are quite challenging and some animals increase their sensory acquisition rate (for review see: (Geva-Sagiv et al., 2015)). For example, animals probing their surrounding through olfaction, increase the sniffing rate when exploring novel objects (Kepecs et al., 2007; Welker, 1964; Wesson et al., 2008). Humans increase the sensory acquisition rate by reducing the rate of eye blinks (Bentivoglio et al., 1997; Shin et al., 2015; Shultz et al., 2011). The present results show that frugivorous bats of the species *C. perspicillata* adapt their sensory acquisition rate in a context-dependent manner, when comparing between challenging and non-challenging conditions. When flying in complex environments, e.g. narrow, cluttered, or noisy areas, the bats increase the acquisition rate by reducing the minimum and median call intervals (Figure 4), by decreasing the call intervals within the call groups (Figure 5) and by increasing the call group size (Figure 5). All adaptations were similar independent from the nature of the complex environment. Thus, the described adaptations may allow the bats to orient collision-free in complex habitats, as it has been suggested for insectivorous bats (Falk et al., 2014; Fawcett et al., 2015; Kothari et al., 2014; Kothari et al., 2018a; Moss et al., 2006; Petrites et al., 2009; Sändig et al., 2014; Surlykke et al., 2009).

Why do bats pattern echolocation calls into groups when orientation becomes demanding? Why do they not simply increase their call rate without grouping the calls? Although there is no direct evidence from *C. perspicillata* that allows to answer these questions, several scenarios seem possible. i) The bats could use the pattern of the call groups to anticipate the correct echo pattern and to associate the echoes to the corresponding calls (Kothari et al., 2018a; Wohlgemuth et al., 2016). For example, a bat emitting a call quartet expects to perceive four echoes with a comparable time pattern as the call quartet. ii) Attentional phenomena often correlate with oscillations of brain activity in the gamma range (higher than 30 Hz; Gregoriou et al., 2009; Gunduz et al., 2011; Sridharan et al., 2011). These oscillations can be imagined as alternating “up” and “down” states of brain activity where “up” stands for high and “down” for low level of attention. The call rate within the call groups lies in the range of 40-50 Hz which might improve stimulus processing by entraining neural activity in the gamma range. Noteworthy, recent neurophysiological data from flying insectivorous bats demonstrated, that the gamma power increases when the bats emit call groups (Kothari et al., 2018b). iii) Another possibility is that each echolocation call of the call group may be spatially directed towards different orientations. Data from the Egyptian fruit-bat *Rousettus aegyptiacus* (Yovel et al., 2010; Yovel et al., 2011) demonstrate that the bats alternate the focus of their sonar beam from left to right and vice versa. This allows a detailed sampling of the distance to surrounding edges and dynamic flight adjustments to avoid sudden collisions. If other bat species (such as *C. perspicillata)* similarly alternate their sonar beam direction is yet to be investigated.

Independent from the reason of patterning echolocation calls into call groups, the behavioral adaptations described in the present study lead to an increased sensory acquisition rate. This allows the bats to gather a detailed representation of the surrounding which might, in turn, help animals to avoid obstacle collision while flying in complex environments.

## Author Contributions

M.J.B. performed experiments. M.J.B. analyzed data. M.J.B. wrote manuscript. M.J.B., J.C.H and M.K. conceived and directed the study. All authors discussed the results and commented on the manuscript.

## Competing interests

The authors declare no competing financial interests.

## Funding

This research was funded by the German Research Foundation (DFG)

